# Statistical signature of subtle behavioural changes in large-scale behavioural assays

**DOI:** 10.1101/2024.05.03.591825

**Authors:** Alexandre Blanc, François Laurent, Alex Barbier–Chebbah, Benjamin T. Cocanougher, Benjamin M.W. Jones, Peter Hague, Marta Zlatic, Rayan Chikhi, Christian L. Vestergaard, Tihana Jovanic, Jean-Baptiste Masson, Chloé Barré

**Author notes:** These authors also contributed equally to this work (Alexandre Blanc, Tihana Jovanic, Jean-Baptiste Masson, Chloé Barré).

## Abstract

The central nervous system can generate various behaviours, including motor responses, which we can observe through video recordings. Recent advancements in genetics, automated behavioural acquisition at scale, and machine learning enable us to link behaviours to their underlying neural mechanisms causally. Moreover, in some animals, such as the *Drosophila* larva, this mapping is possible at unprecedented scales of millions of animals and single neurons, allowing us to identify the neural circuits generating particular behaviours.

These high-throughput screening efforts are invaluable, linking the activation or suppression of specific neurons to behavioural patterns in millions of animals. This provides a rich dataset to explore how diverse nervous system responses can be to the same stimuli. However, challenges remain in identifying subtle behaviours from these large datasets, including immediate and delayed responses to neural activation or suppression, and understanding these behaviours on a large scale. We introduce several statistically robust methods for analyzing behavioural data in response to these challenges: 1) A generative physical model that regularizes the inference of larval shapes across the entire dataset. 2) An unsupervised kernel-based method for statistical testing in learned behavioural spaces aimed at detecting subtle deviations in behaviour. 3) A generative model for larval behavioural sequences, providing a benchmark for identifying complex behavioural changes. 4) A comprehensive analysis technique using suffix trees to categorize genetic lines into clusters based on common action sequences. We showcase these methodologies through a behavioural screen focused on responses to an air puff, analyzing data from 280,716 larvae across 568 genetic lines.

**Author Summary:** There is a significant gap in understanding between the architecture of neural circuits and the mechanisms of action selection and behaviour generation.*Drosophila* larvae have emerged as an ideal platform for simultaneously probing behaviour and the underlying neuronal computation [1]. Modern genetic tools allow efficient activation or silencing of individual and small groups of neurons. Combining these techniques with standardized stimuli over thousands of individuals makes it possible to relate neurons to behaviour causally. However, extracting these relationships from massive and noisy recordings requires the development of new statistically robust approaches. We introduce a suite of statistical methods that utilize individual behavioural data and the overarching structure of the behavioural screen to deduce subtle behavioural changes from raw data. Given our study’s extensive number of larvae, addressing and preempting potential challenges in body shape recognition is critical for enhancing behaviour detection. To this end, we have adopted a physics-informed inference model. Our first group of techniques enables robust statistical analysis within a learned continuous behaviour latent space, facilitating the detection of subtle behavioural shifts relative to reference genetic lines. A second array of methods probes for subtle variations in action sequences by comparing them to a bespoke generative model. Together, these strategies have enabled us to construct representations of behavioural patterns specific to a lineage and identify a roster of ”hit” neurons with the potential to influence behaviour subtly.

## Introduction

Animals integrate external sensory input and their internal states to generate suitable motor responses. This involves different areas of the nervous system, ranging from areas underlying sensory processing and higher-order processing to those governing decision making and motor control. Furthermore, animals frequently respond to stimuli with a sequence of actions requiring precise control of transitions between individual actions. Different animals may react differently to the same stimulus, and the same animal can respond variably to repeated stimuli. This probabilistic nature of responses implies complexity and stochasticity in the behavioural choice mechanisms. The neurobiological interactions among neurons that regulate the trade-off between action stability and variability and control transitions between actions remain only partially understood.

Identifying the neural substrates that are responsible for behaviour generation and selection within the nervous system is crucial. Historically, this task has been challenging due to simultaneously manipulating neuron groups while capturing the corresponding behaviours and the statistical complexities involved in causally linking behavioural sequences across multiple time scales to neuronal manipulations.

The past decade has witnessed significant advancements in connecting behaviours with neural computations. Notably, data-driven neuron-behavior mappings have been established for *Drosophila melanogaster* in both its adult [2] and larval stages [3]. *D. melanogaster* presents an ideal model for such studies due to its sufficiently complex yet accessible nervous system, comprising roughly 10,000 neurons in larvae and 130,000 neurons in adults. The complete synaptic connectomes for larval (full CNS connectome) and adult (brain) *D. melanogaster* are now fully mapped [4–6], providing detailed diagrams of neuronal connections. Additionally, the *D. melanogaster* genome has been extensively characterized, and the development of thousands of GAL4 lines facilitates precise genetic manipulation [7, 8], nearly down to the level of individual neurons.

The semi-transparent cuticle of the larva enables the application of optogenetic techniques to selectively and reproducibly activate or inactivate neurons during behaviour across the entire nervous system [3, 9]. Techniques such as the targeted genetic expression of tetanus neurotoxin (TNT) can also disrupt synaptic transmission in small or individual neuron groups. High-throughput tracking with real-time segmentation capabilities allows for recording hundreds of thousands of larvae, with individual neurons or neuron groups being selectively activated or silenced, constitutively or reversibly [10].

Advances in machine learning [11, 12] have recently complemented automated behavioural analyses, and supervised [13–20] and unsupervised methods [21–29] have been introduced alongside image feature-based approaches to identify behaviours. Some methods can be applied broadly to various experiments after an annotation phase, like DeepLabCut [13], or, as with some unsupervised approaches, others are more specialised and apply to one animal in a specified behavioural paradigm [30, 31]. Supervised techniques aim to define behaviours based on external expertise, while unsupervised ones seek to have them naturally emerge, later undergoing post-hoc validation by experts. Overall, the success of these methods depends on the definition of behaviours, the amount of accessible data and its standardisation, and the variability expected under the experimental protocol. Usually, these frameworks link sensory stimuli or targeted neural activation to their behavioural output and are associated with statistical testing to detect significant events.

The primary challenges posed by the behavioural recordings of larvae are linked to the significant deformability of their bodies, the low resolution of images, which is imposed to allow large-scale screening, the multi-temporal scales of their behavioural dynamics, and the vast diversity of larval morphological characteristics across populations of several hundred thousand animals. In spite of these complications, both unsupervised [3, 24] and supervised [32–34] approaches have been successfully applied, albeit with known limitations. In supervised approaches, ambiguityies in larva behaviour prevent full consensus on behavioural ground truth. New experiments suggest that additional actions may be required to properly describe larval behaviour, such as its C-shape behavior before rolling [35, 36]. Furthermore, the diversity of larvae lengths, speeds, variations in the recording time of the larva, and the inherent deformability of the larva body induce challenges in estimating behaviour classification errors. Finally, these ambiguities shift the identification of neurons of interest, i.e., neurons able to modify the larva behaviour, towards the ones inducing large behavioural deviations.

In this paper, we develop new statistical tests allowing the detection of neurons inducing subtle changes in behaviour. To ensure the robustness of such finer analyses, we first introduce a physics-informed Bayesian approach to regularise the recorded shapes of the larvae. We then introduce two statistical approaches to provide a global analysis of the larva behavioural screen and identify neurons able to induce subtle variations in local behaviour or in the higher-order statistics of sequences of actions. We apply these approaches on an entire behavioural screen and demonstrate the ability of both approaches to detect neurons or group of neurons able to induce subtle behavioural changes. We leverage our new approaches to provide compact representation of lines behvioural phenotypes and

## Materials and methods

### *Drosophila melanogaster* stocks

The screen consisted of 569 GAL4 lines, as listed in Table 1. These lines were from the Rubin collection lines (available from Bloomington stock centre) listed in S1 Data file, each of which is associated with an image of the neuronal expression pattern shown at flweb.janelia.org/cgi-bin/flew.cgi. In addition, we used the insertion site stocks, w;attP2 [7], OK107GAL4, 19-12-GAL4, NompC [37], and iav-GAL4 [38]. We used the progeny larvae from the insertion site stock, w;;attp2, crossed to the appropriate effector (UAS-TNT-e (II)) for silencing. The w;;attP2 were selected because they have the same genetic background as the GAL4 tested in the screen. We used the following effector stocks: UAS-TNT-e [39] and pJFRC12-10XUAS-IVSmyr::GFP (Bloomington stock number: 32197).

### Behavioural apparatus, experiments and screen design

#### Apparatus

The setup was fully described previously [10, 34] (see Fig. 1). Briefly, it consists of a video camera for monitoring larvae, a ring light illuminator, and custom hardware modules for generating air puffs, controlled through the multi-worm tracker (MWT) software [9, 40].

**Fig 1.**
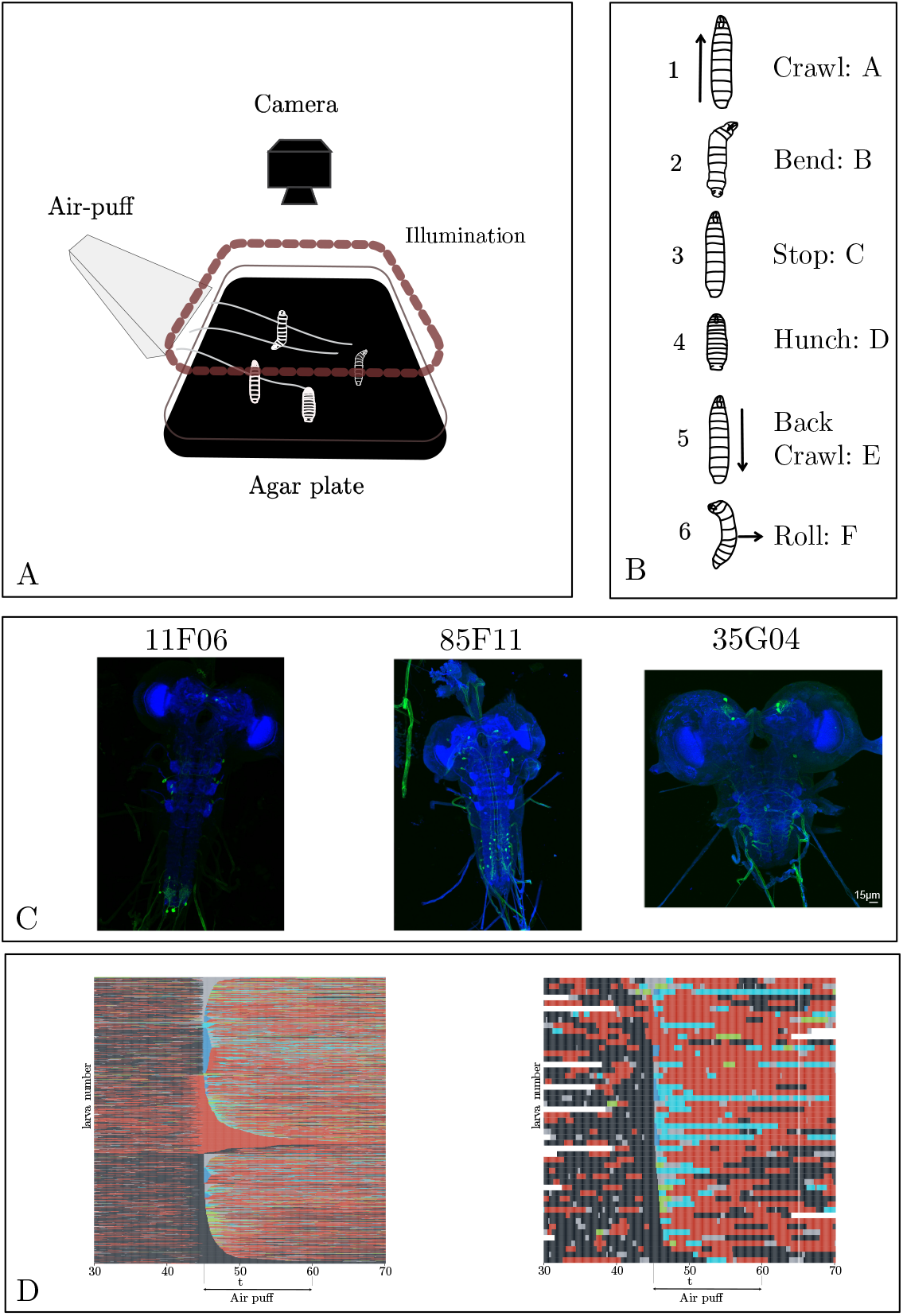
(A) Behavioural set-up. The larvae move freely on an agar plate, and their movement is recorded with an infrared camera equipped with a high-throughput closed-loop tracker. The stimuli were an air puff (or illumination for training data). (B) The six stereotypical actions [9, 10] associated with the larva for this experimental paradigm. (C) Example of Neuronal expression patterns in three example lines: 11F06, 85F22, and 35G04. (D) Ethogram of larva behaviour in response to an air-puff at 45s based on automated behavioural detection. Each line corresponds to one larva, with on the left the Control line, attP2, and on the right R35G04. Each colour corresponds to one of the six actions: black: crawl, red: bend, blue: stop, deep blue: hunch, and cyan: back (no rolls were observed in these lines).

#### Behavioural experiments

The experiments are fully described in [10]. Briefly, they started with collecting embryos for 8–16 hours at 25°C with 65% humidity. Larvae were raised at 25°C with normal cornmeal food. Foraging 3rd instar larvae were used (larvae reared 72–84 hours or for three days at 25°C). Before experiments, larvae were separated from food using 10% sucrose, scooped with a paintbrush into a sieve and washed with water. The substrate for behavioural experiments was a 3% Bacto agar gel in 25 625 cm^2^ square plastic dishes. Batches of approximately 50 to 100 larvae were imaged in each behavioural assay. The larvae were left to crawl freely on an agar plate for 44 seconds before the stimulus delivery. The air puff was delivered at the 45th second and applied for 38 seconds. Two different stimulus intensities were considered, one at a high intensity of 6 m/s and the other at a lower intensity of 3 m/s. In this paper, when a result is stated without indicating a specific intensity, it must be understood that it was obtained with the higher 6 m/s.

#### Screen design

The screen consisted of recordings of the behaviour of 568 GAL4 lines from the Rubin GAL4 collection, where we constitutively silenced small subsets of neurons and individual neurons using tetanus toxin [34, 39]. We selected these lines from the entire collection for sparse expression in the brain and ventral nerve cord of the larval CNS and expression in the sensory neurons. Of the 569 lines tested here, several neuronal lines were not part of the Rubin collection: we added 19–12-GAL4 and NompC-GAL4 for sensory neurons and OK107GAL4 for the mushroom body. We screened each GAL4 line in the air-puff assay described above. This article used no activation method (optogenetic or other) since we used constitutive silencing.

#### Behavioural dictionary

Six stereotypical actions are commonly used to constitute the behavioral dictionary of the larva (Fig. 1B): A: crawl, B: bend (all turning actions), C: stop (not moving), D: hunch (fast retraction of the head) E: back crawl (crawling backwards), and F: roll (defensive manoeuvre consisting in sliding laterally). We use the letters A–F in plots and tables for brevity. Where these actions were needed for the analysis, we inferred them using a combination of supervised and unsupervised machine-learning techniques introduced in [34].

### Physics-informed regularization of larva shape

We conducted large-scale imaging by recording larvae with a wide-field view, allowing us to analyze up to 100 larvae per plate. This approach, while time efficient, resulted in images of lower resolution. Additionally, the vast scale of our experiments meant that many larvae were not perfectly dried, leading to abnormal contour shapes. Impurities in the agar further contributed to these irregularities, as illustrated in Figure 2. Such contour abnormalities risk leading to misclassification of larval behaviour, potentially introducing bias into subsequent statistical analyses. To address this, we developed a regularisation procedure based on physics-informed Bayesian inference [41], which ensures accurate representation of larval shapes.

**Fig 2.**
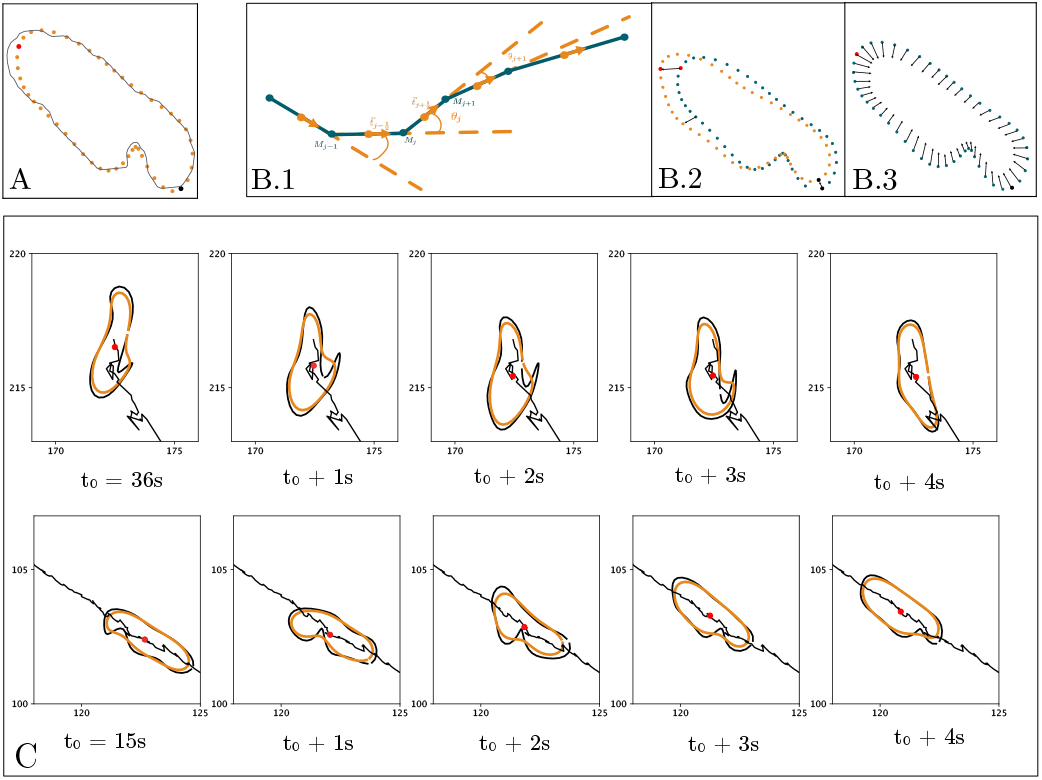
(A) Tracked contour of a noisy outline of a larva in grey and regularised contour in orange with the head in red and the tail in black. (B) 1. Zoom on six points of the larva contour, the contour is materialized by vectors between these points. The *j*th point is named *M*_*j*_, its tangent vector 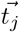, and the curvature in the point *θ*_*j*_. 2. Two outlines of a larva at time *t* and *t* + *dt*, the vectors show the movement of two points during the time-lapse *dt*. 3. Arrows showing the movement of the contour points as the surface energy is decreased. (C) Results after running the algorithm for two different larvae at four different time steps with the old outline in black and the new one in orange. The trajectory of the larva is drawn in black, and the centre of mass is marked by a red dot (see also Supplementary Video 1).

#### Preprocessing

MWT extracts contours with a variable number of points depending on the larvae’s size in each frame. We denote this contour by *f* (*i*) = (*x*(*i*), *y*(*i*)) for *i* ∈ *{*1, 2, …, *N*_tracking_*}* with *N*_tracking_ the number of points on the contour. We regularized the shape by fixing the number of contour points to *N* = 50 coupled with a low-pass filtering. In paticular, we generated the contours by retaining the *K* lowest modes of the Fourier decomposition [42] of the recorded contour (Fig. 2A),

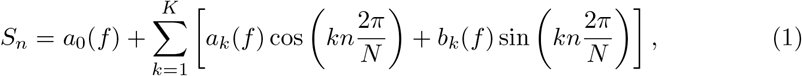

with *a*_*k*_ and *b*_*k*_ the Fourier coefficients,

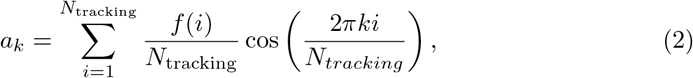

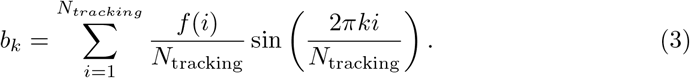

We reconstructed the shape with the *K* = 7 lowest harmonics, a number chosen empirically to prevent discontinuities [42]. This first reconstruction ensured screen-scale regularisation of larvae contours regardless of their variability in size and shape.

#### Simplified physics model of larva

We designed a minimal, effective 2D physics model to approximate the dynamical shape of the larva. It models the larva as an elastic contour with an active membrane energy. The total energy of the larva is the sum of the kinetic energy, the surface energy, and the bending energy: *E* = *E*_*k*_ + *E*_*S*_ + *E*_*b*_ (Fig. 2B.2). The kinetic energy is given by

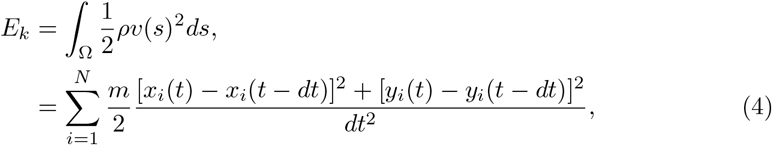

where Ω is the surface of the contour, *ρ* is the surface density, *v*(*s*) is the speed of the contour in the point *s, m* is the total mass of the membrane, and *dt* is the time lapse between images. The surface energy is

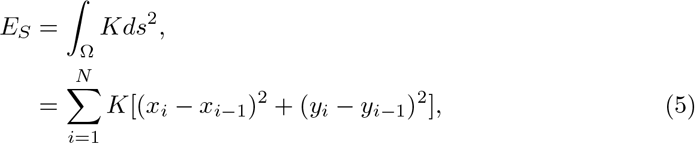

with *K* the elastic modulus. Finally, the bending energy reads as

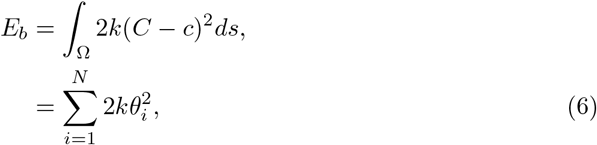

where *k* is the bending modulus, *C* is the mean curvature over the entire contour, and *c* is the spontaneous curvature defined by 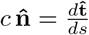 with 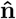 the unit normal vector and 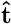 the unit tangent vector of the contour in curvilinear coordinates. For a discrete point, this curvature equals *c*_*i*_ = *θ*_*i*_ (see Fig. 2B.1), and we will set the mean curvature to 0.

#### Inference

We used Bayesian inference to infer the larva’s regularised shape 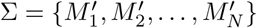. Its posterior distribution is given by

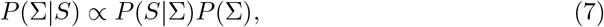

where *P* (*S* | Σ) us the likelihood of the model and *P* (Σ) is the prior which regularises the inference by incorporating our physical model. It is given by

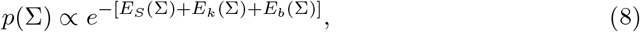

up to a normalising constant that does not influence the inference. The likelihood enforces the proximity between the recorded contour and the inferred one according to a quadratic loss,

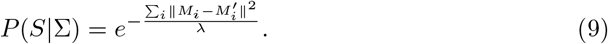

We set 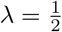 with no loss in generality since the absolute scale of the energy does not impact the inference.

The log-posterior distribution thus reads

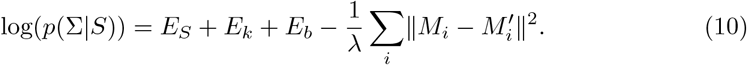

The model’s hyperparameters (*m, k*, and *K*) were set to give the three terms of the energy similar weights. The larva’s mass was set to *m* = 1, the curvature coefficient to *k* = 1, and the elastic modulus to *K* = 5. In the numerical implementation, we corrected spurious high curvature anomalies by capping the energy using 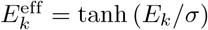 with *σ* = 100. We used stochastic gradient descent [43] to infer the maximum a posteriori (MAP) regularised contour Σ. We show in Figure 2C inferred contours in 2 examples displaying significant anomalies. Note that while we rely solely on the MAP of the shape in downstream analysis and not on the full posterior distribution, it is accessible using Markov Chain Monte Carlo [43] sampling if necessary.

### A continuous self-supervised representation of behaviour

We developed a continuous representation of the behaviour based on self-supervised learning (SSL) to alleviate the need for a predefined behavioural dictionary to characterize larvae actions. SSL is a general paradigm [44–50] in which a model is trained using auxiliary objectives to improve the performance for downstream tasks. The auxiliary objectives are generally constructed from the data themselves, thus requiring no external labelling. Here, we present our implementation based on larva positional prediction of the regularized shapes inferred as described above (Physics-informed regularization of larva shape).

#### Architecture and training of the neural network

We used an autoencoder architecture comprising an encoder and a decoder, mapping input and output data, as illustrated in Figure 3A. The encoder takes as input a sample, *X*_*t*_ and maps it to a learned latent space, producing the latent representation of the sample. The decoder takes as input a latent representation and generates a reconstruction of the sample, 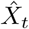, in the data space. We augment the objective by requiring the decoder to also predict the preceding and following samples, *X*_*t*−1_ and *X*_*t*+1_, forcing the neural network to encode the temporal continuity of the larvae’s motion.

**Fig 3.**
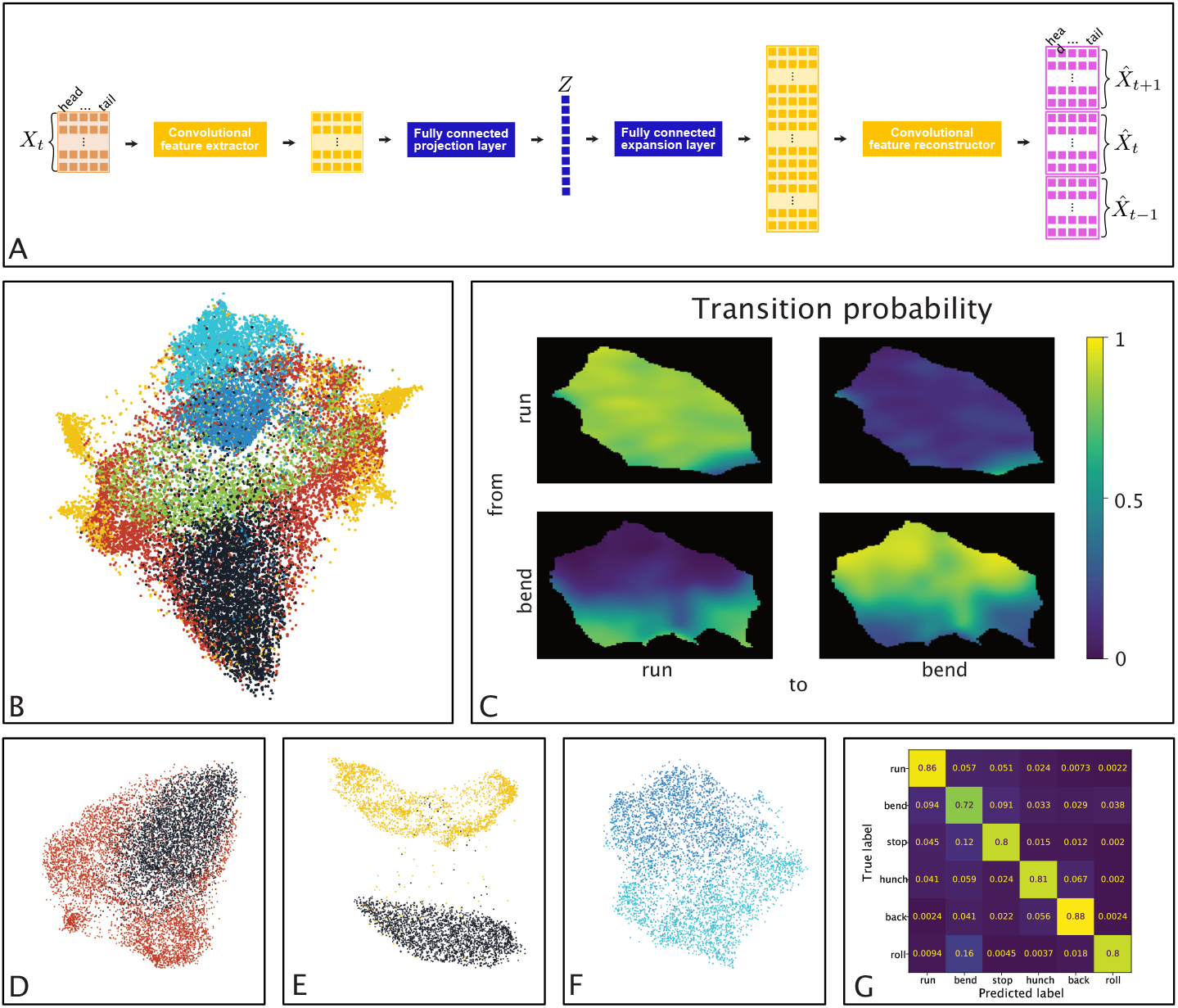
(A) Architecture of the self-supervised predictive autoencoder. The encoder consists of multiple convolutions with ReLU activations alternating between the spatial and temporal axis of the data, followed by a fully connected linear layer. The decoder consists of an upsampling linear layer matching the internal representation to the desired shape, followed by alternating convolutions with ReLU activations. (B) Visualization of the latent space. The 10D latent space is projected in 2D using UMAP [51]. The colours correspond to the discrete behaviour dictionary (black: crawl, red: bend, green: stop, blue: hunch, cyan: back, and yellow: roll) (C) Transition probability from one discrete state to the other as a function of the position in the latent space: here, between run and bend. (D–F) Highlights of the behaviour geometry in the latent space (represented in 2D using UMAP). In D run vs. bend, in E run vs. roll, and in F hunch vs. back. (G) Cross-validated confusion matrix of random forest classifiers using the latent representation to infer the usual discrete behaviour dictionary.

We used an epoch approach to encode the dynamics of the larva with a predefined time interval of length *τ*. Unless indicated otherwise, we used *τ* = 2s. In principle, *τ* could range from very short durations (a few hundred ms), capturing primitive muscular responses, to longer durations (5-10s), capturing entire sequences of actions. The choice *τ* = 2s was informed by previous behaviour analysis of larvae [9, 10, 34, 52] and chosen to ensure capturing transitions between different actions. The autoencoder was tasked (see Fig 3A) to encode the coordinates of the larva within the present epoch and to predict the coordinates for the immediate future and past epoch (of the same duration *τ*). The autoencoder was trained to minimize an *L*^2^ objective on the reconstructed sequences. The autoencoder performs both 2D-convolutions and 1-D convolutions, acting alternately on the spatial and temporal coordinates. Hyperparameters, architecture, and source code are provided at `dedicated.github`.

#### Datasets for training and testing

Tracking data were initially generated using MWT [40] as described above. We post-processed them using the pipeline introduced in [9], limiting the representation of a larva to 5 points along its anteroposterior axis [9] (tail, lower neck, neck, upper neck, head) to allow the representation to be used with very low-resolution imaging of larvae (as in [53]).

We assigned one of the six following discrete behavioural categories to each time point of the larva: *run, bend, stop, hunch, back* and *roll* using the pipeline of [34]. To promote generalisation across lines and robustness regarding different morphologies, larva coordinates were normalized such that the line average of the larva length before sensory stimuli was equal to one. Furthermore, we centred and aligned the larva so that their average position was zero and their average orientation along the x-axis, towards negative *x*. Data were sampled from experiments in [3] and in [1, 10, 34, 54]

Natural behaviour statistics are deeply imbalanced. Before the sensory input signal, the animals are freely moving with roughly 70% run and 30% bend with occasional stops. The sensory stimulus can generate behaviour that would not be evoked without stimulation. All data pooled together, regardless of experimental protocols, exhibited the following statistics with Run: 50.28%, Bend: 39.35%, Stop: 6.43%, Hunch: 0.81%, Back: 3.06%, and Roll: 0.07%. Lexical approaches such as the one from [22], while very efficient in analysing animal behaviour in natural settings, have challenges with such a level of imbalance.

We used an inductive bias to train the autoencoders. Training data consisted of 100 000 samples, 10% of which were held out for validation, with 25% runs, 25% bends, and 12,5% of each of the other four behaviours.

### Genotype-level analysis

#### Genotype representation

To detect genotypes of interest (commonly called *hits*), we employ a non-parametric statistical test within the latent behaviour space (learned as described in the section A continuous self-supervised representation of behaviour). While the testing relies on the learned behavioural space, we emphasize that other architectures and objectives (such as [23]) may be used if they provide a sufficiently robust description of the behaviour.

Following the stimulus, we immediately embedded the *τ* -long windows of behaviour, resulting in a sample of behavioural responses represented as points in the latent space. One larva’s behavioural dynamics becomes a singular point within the latent space. A genotype’s experimental behavioural dynamics, evaluated, for example, on 1000 larvae, becomes a distribution of 1000 points inside the latent space. We estimated the underlying distribution using a Gaussian kernel. Therefore, the phenotypic characterisation of a genotype reduces to a probability distribution in the learned latent space, so we reduce the comparison of two genotypes to a comparison of two probability distributions in a low-dimensional space (Fig. 4A).

**Fig 4.**
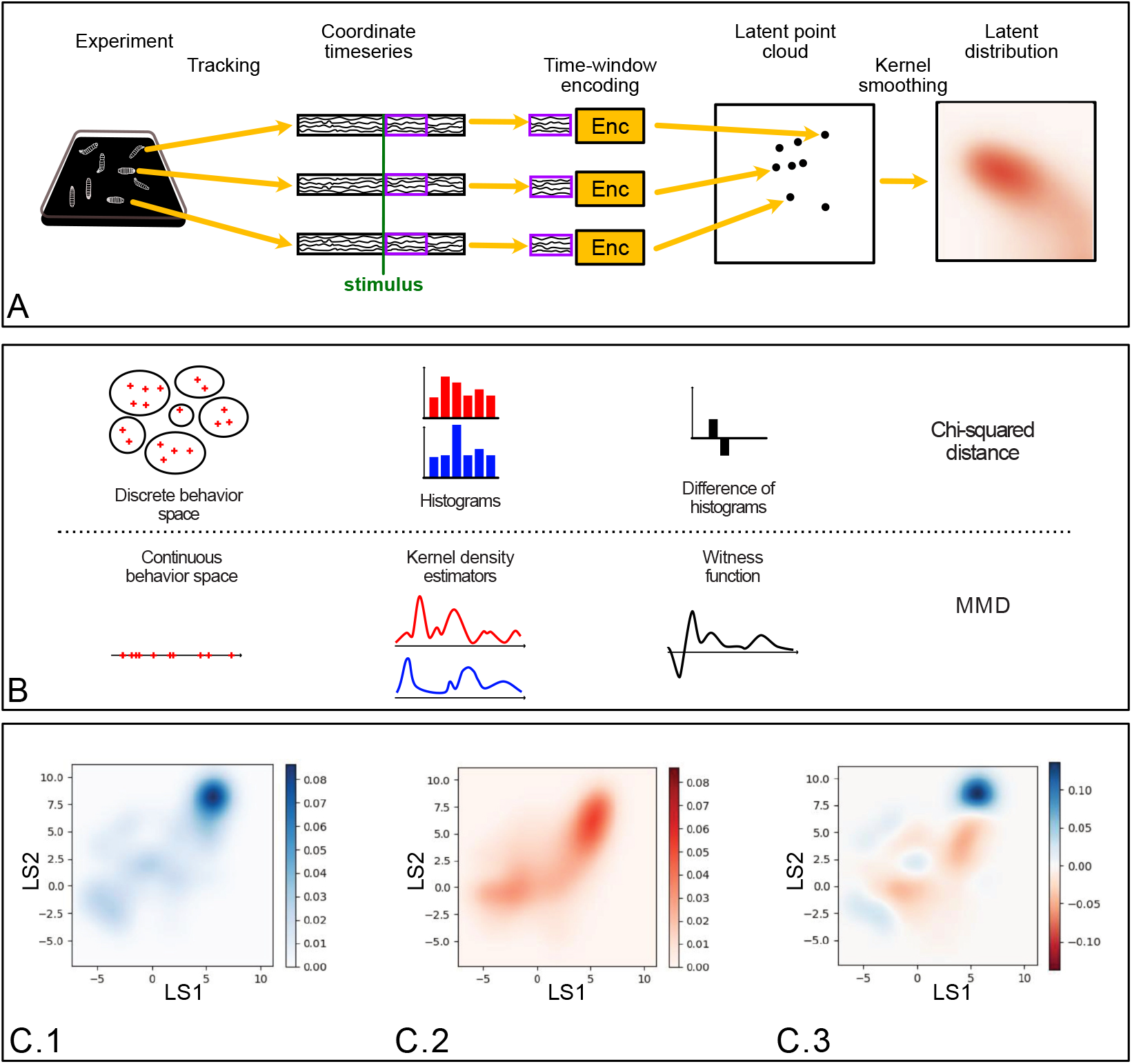
(A) Illustration of our phenotyping modelling strategy for each genotype. From left to right: The behaviour evolution on the experimental setup reduced to the five tracked points of the larva, the extraction of a temporal window (shown in purple on the ethogram as an illustration) usually after the onset of the stimuli (shown as a vertical green line), projection of the temporal window on the latent space using the encoder shown in Fig. 3 and reduced here to a yellow box, each point in the latent space corresponds to one larva behaviour during the selected time window, the phenotype of the genotype is the distribution of all the points in the latent space regularised by a Gaussian kernel. (B) Illustration of the correspondence between statistical testing procedures based on discrete behaviour categories with chi-squared tests and testing procedures based on continuous behaviour with MMD. (C) Latent distributions of behaviour (regularized by a Gaussian kernel): (C.1) of the reference line *attP* 2 and (C.2) of the line 10*A*11. (C.3) Witness function between these two latent distributions, highlighting main behavioural differences between the lines.

#### Kernel-based statistical testing

We used the maximum mean discrepancy [55] (MMD) to measure the distance between distributions (Fig. 4B), which is efficient in detecting subtle differences between datasets [56]. MMD was developed to perform non-parametric statistical testing between two sets of independent observations in a metric space *Ƶ* (here the latent space *Ƶ* = ℝ^10^). We denote by *X* = *{x*_1_, …, *x*_*m*_*}* the first set, drawn from the distribution *p*, and by *Y* = {*y*_1_, …, *y*_*n*_} the second, drawn from *q*. The goal is to test if *p* = *q*, i.e., we seek to reject the null hypothesis that the two genotypes have similar behavioural responses.

The MMD between two probability measures *p* and *q* is defined as

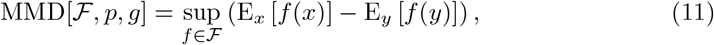

where *ℱ* is a class of functions from *Ƶ* to ℝ, and E_*x*_ and E_*y*_ denote expectation w.r.t. *p* and *q*, respectively.

When the function class is the unit ball in a reproducing kernel Hilbert space *ℋ*, the square of the MMD can be estimated directly from data samples [57]. We estimated the squared MMD between *X* and *Y* using the unbiased estimator given by

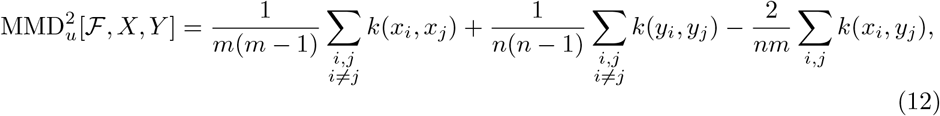

where *k* denotes the kernel operator, here a Gaussian kernel given by 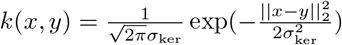. The bandwidth *σ*_ker_ was calibrated using the median of the pairwise distances in the latent space of samples corresponding to the reference line following [55].

The MMD framework provides explainability of the statistical test by enabling identification of the variables that exhibit the greatest difference between datasets [57, 58]. It defines a particular function over the vector space that supports the distributions, called the witness function that highlights regions where large deviations occur, Fig. 4C. These regions can be analysed further to identify the behavioural features associated with them.

### Probabilistic generative model of action sequences

In addition to the dictionary-free approach for behavioural analysis (see the section A continuous self-supervised representation of behaviour) that compares various larva genotype responses to the air-puff stimuli to the reference genotype, we developed a structured probabilistic approach to probe higher-order behavioural patterns that directly influence action sequences. This approach compares each genotype to a constrained generative model with a behaviour dictionary instead of relying on direct comparison to a reference genotype.

The sequences of actions are modelled using a time-varying continuous-time Markov chain, built upon simple probabilistic basis functions that draw inspiration directly from bacterial chemotaxis (see e.g. [59]). The model is parametrised by the average duration of each action, the action’s amplitude (either the maximum asymmetry factor (an experimentally robust proxy to the bending angle) or the velocity of the action), and the transition probabilities between successive actions. All parameters are allowed to vary temporally. We account for this time variation by using piecewise constant parameters in a *δt* = 1 s time windows.

We initialise the state of the larva following the stationary distribution of actions, *p*(*i, t*_0_). At a given point in time *t* (including *t* = *t*_0_), the duration Δ*t* of an action is drawn from a Poisson distribution,

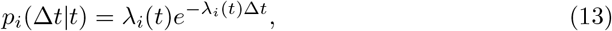

with 1/λ_*i*_(t) the mean time spent in behaviour *i* at time *t*. The action’s amplitude is allowed to depend on Δ*t*. For asymmetric actions (i.e., bend, hunch, and roll) the amplitude is quantified by the ”asymmetry” factor *A*_*s*_, while for the other actions (i.e., run, back, and hunch) it is quantified by the velocity *v*. The asymmetry is defined by 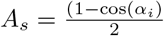, with *α*_*i*_ the angle between the segment formed by the centre and the head of the larva and the segment formed by the centre and the tail of the larva. The asymmetry can only take values in the [−1, 1] range. We approximate the amplitude distributions using a kernel density estimation with a mixture of Gaussian kernels with uniform weights,

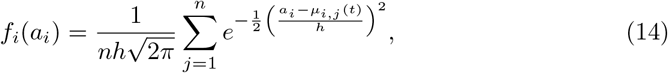

where *a*_*i*_ is the amplitude of the velocity or the asymmetry of the behaviour *i, µ*_*i*,*j*_(*t*) is the mean of *j*th component of the mixture, 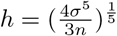 is the bandwidth of the kernel, and *σ* the standard deviation of the amplitudes inferred from data. We set *n* = 10, which empirically leads to a good trade off between bias and variance. Thus, the distribution is described by a 10-dimensional vector *µ*_*i*_(*t*) = (*µ*_*i*,1_(*t*)(*t*), …, *µ*_*i*,10_(*t*)). Finally, a new action is chosen according to a first-order Markov chain parametrised by the transition matrix *T* (*t*). Since we explicitly model action duration, self-transitions are not possible and the diagonal elements of *T* (*t*) are zero.

The full set of parameters to infer is (**Λ, M, T**). Here **Λ** = *{λ*_*i*_(*t*)*}* is the expected inverse durations of each behaviour during each second. **M** = {*µ*_*i*_(*t*, Δ*t*) } is the features for each action during each second knowing the duration of the action, and with *µ*_*i*_(*t*, Δ*t*) = (*µ*_*i*,1_(*t*)(*t*), …, *µ*_*i*,10_(*t*)). **T** = {*T* (*t*) } is the set of transition matrices over time, also changing each second.

The model’s parameters are learnt from experimental data using Bayesian inference (see Eq. 7). The likelihood for one larva’s behaviour sequence can be written as

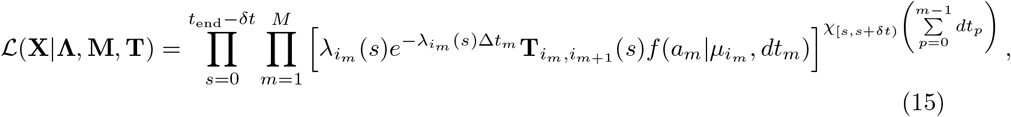

where **X** = {(*i*_1_, *a*_1_, Δ*t*_1_), …, (*i*_*M*_, *a*_*M*_, *δt*_*M*_)} is the sequence of the larva’s actions, *t*_end_ is the duration of the recording, *χ*_[*a*,*b*)_(*x*) is indicator function for *x* being in the interval [*a, b*), and *i*_*m*_ is the *m*th action, with Δ*t*_*m*_ its duration and *a*_*m*_ is its amplitude. The products

We regularise the inference using the following priors on the temporal variation of the parameters:

- a prior enforcing the normalization of the transition matrix, 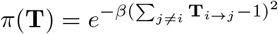;
- a prior reinforcing a smooth temporal variation of **Λ**, 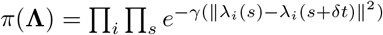;
- s prior reinforcing a smooth temporal variation of **M**(**t**), 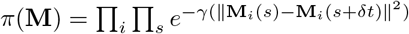,

The maximum a posteriori values of the parameters are inferred by minimizing the following cost function

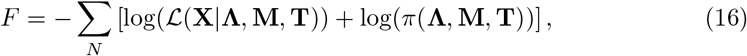

where *π*(**Λ, M, T**) = *π*(**Λ**)*π*(**M**)*π*(**T**) and *N* is the number of larvae. We minimize this function with a direct gradient descent algorithm on the entire set of behavior sequences of the larvae of a single genotype. We visually represent the model in Figure 5A.

**Fig 5.**
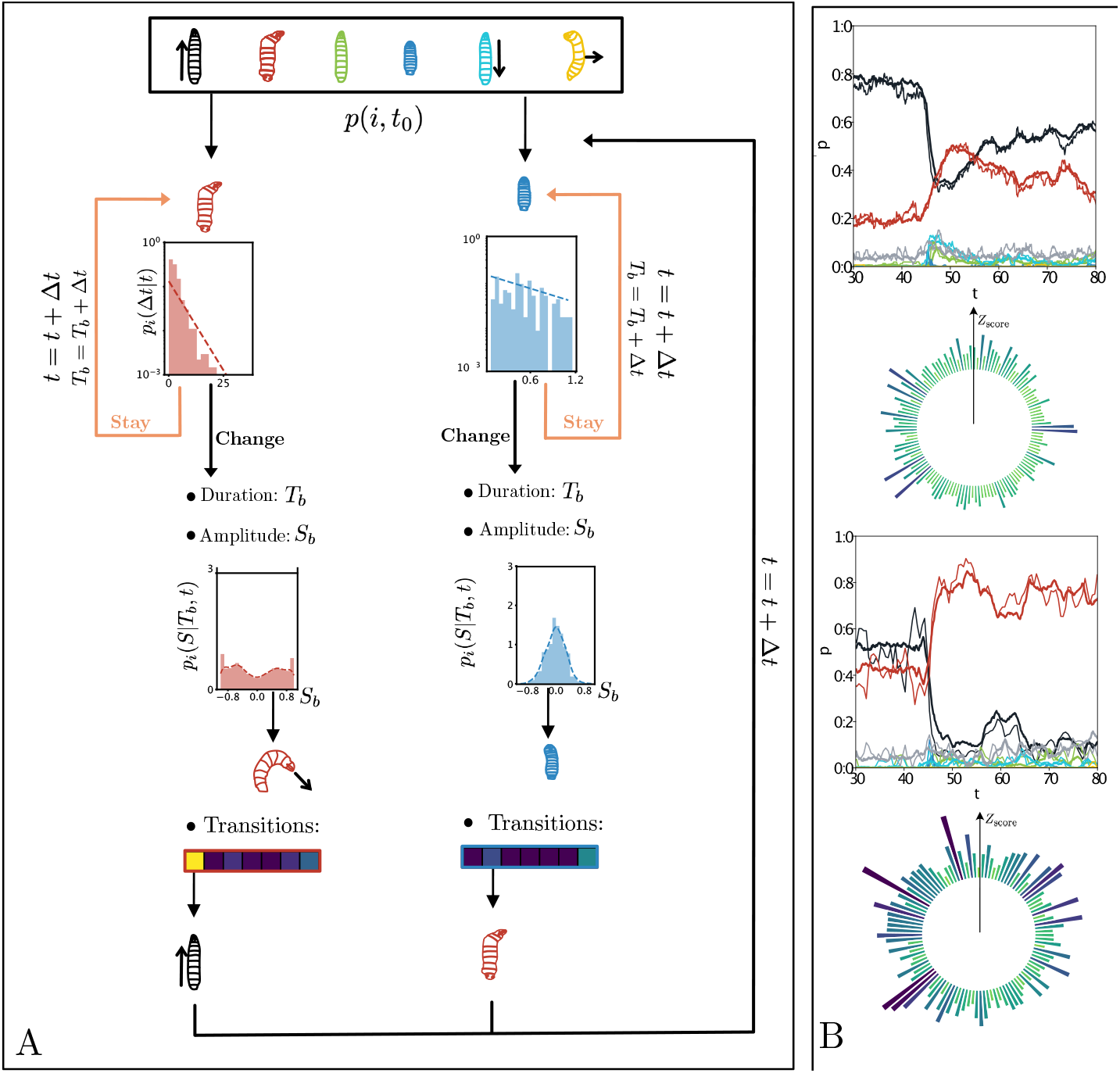
(A) Graphical representation of the probabilistic generative model showing the temporally inhomogeneous Poisson model *p*_*i*_(Δ*t*|*t*), the distribution of action amplitudes *p*_*i*_(*S* | *T*_*b*_, *t*), and transition probabilities to the other actions. (B) Characterisation of behavioural responses to an air puff with the prediction of the generative model for two lines. On top: time evolution of the larva’s actions; thin lines are the experimental recording, and thick lines are the generative model; on the bottom, a circular plot of the z-scores between the action sequences of the generative model and the experimental recordings. Darker blue colours indicate larger values. The two lines are *R*41*D*01 on top and *R*38*H*09 on the bottom.

After inference, we generate artificial behavioural sequences from the inferred parameters from our model using Monte Carlo sampling of the posterior distribution and generate new artificial sequences using the procedure outlined in 22.

We evaluated the model’s goodness of fit using the MMD and compared generated sequences and real sequences of behaviours. For each line, we took groups of 100 random larvae, for which we calculated the probabilities of sequence occurrence. We obtain a distance for each line corresponding to the differences between the models and the experiments. We provide the distances in Supplementary Table 2.

We added to the global scoring performed by the MMD with sequence-based scoring, allowing direct comparison of the probability of a defined sequence under the fitted model and its experimental frequency. We used the z-score 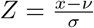 with *x* the probability of the sequence under the model, *v* the frequency of the sequence in experimental data and *σ* a bootstrap estimation of the standard deviation of the sample frequency under the generative model. We limited the analysis to sequences of 3 actions to maintain statistical significance for most lines in the screen. Figure 5B shows an example of Z score for all sequences for two different lines. Although the model reproduces the evolution of probabilities over time, some sequences on line 38H09 are poorly described, as evidenced by their large Z score (the values of the Z scores per sequence for certain lines are noted in Table 3.1 and Table 3.2 (during the stimulus, and during all time)).

### Clustering behavioural sequences from suffix tree representations

The total number of actions performed during this screen is roughly 1.3 million. The scale and diversity of recorded behaviour can be exploited to identify subtle structures in behavioural sequences by analysing the screen as a general ensemble. We used a suffix tree representation to explore the entire screen structure and the genetic lines’ organisation. We constructed the suffix tree with Ukkonen’s algorithm [60, 61] on all sequences of behavior of all larvae across all genetic lines. We here consider only the sequence of categorical behavioural actions, regardless of their durations. Each sequence is added to the suffix tree (see Fig. 6A for a illustrative example of a tree built from just 3 different larvae). The size of the tree grows quadratically with the length of the sequence, and the proportion of sequences common to several lines decreases accordingly. Thus to avoid too long sequences (which would decrease statistical power due to their multiplicity), and to focus on the biologically relevant immediate response behaviours, we only consider behaviors occurring during the first 5 seconds after the onset of the stimulus. (This has the additional benefit of limiting the computational burden.)

**Fig 6.**
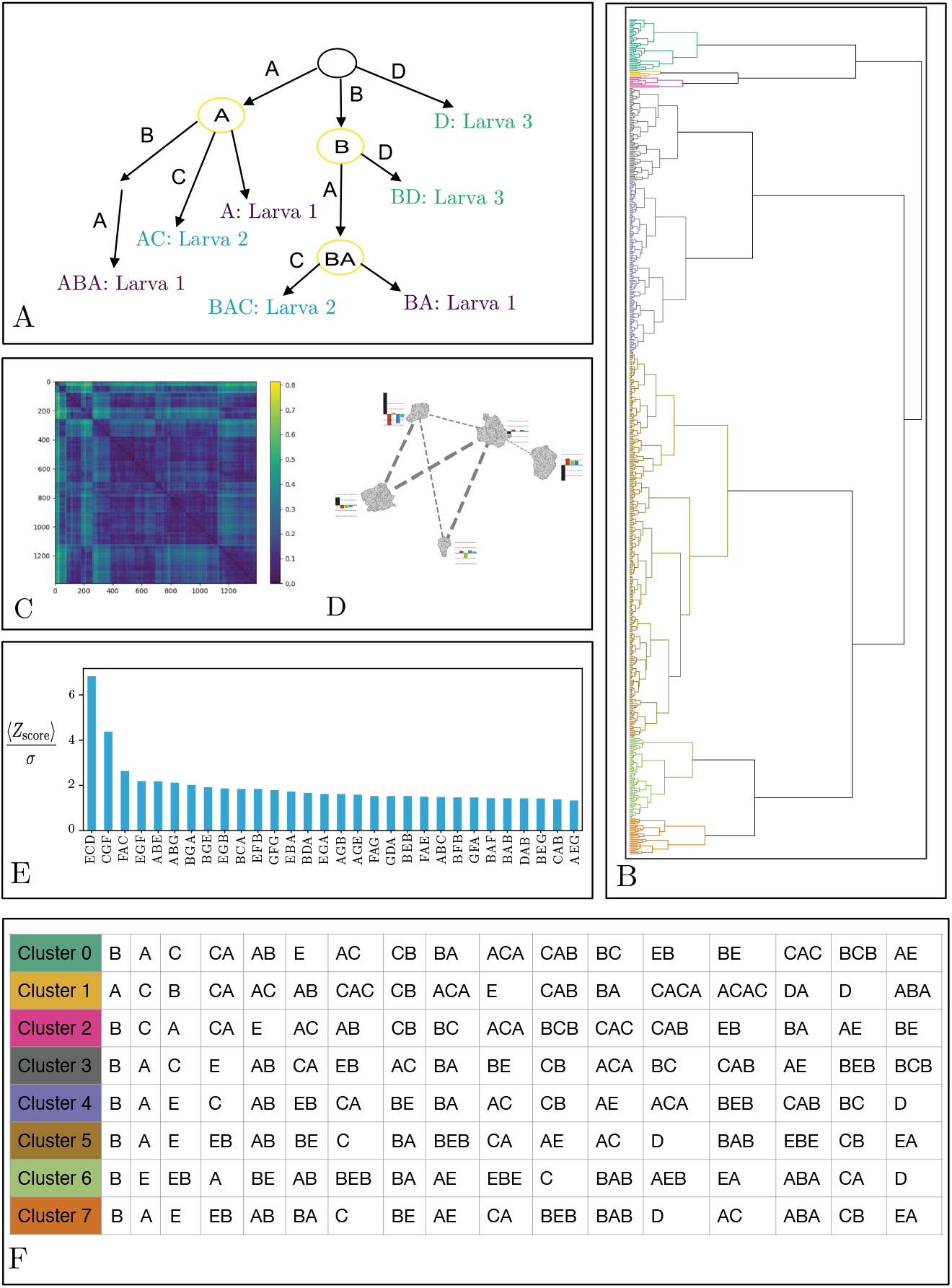
A. Schema of a suffix tree for three larvae performing three different sequences. Larva 1: ABA, Larva 2: BAC, Larva 3: BD, the seven paths from the root to the leaves correspond to the seven suffixes: A, BA, ABA, AC, BAC, D and BD. Each node shared by at least two larvae is shown in circles: A, B and BA. B. Hierarchical clustering based on the cosine similarity between the suffix tree vectors of each genetic line. Each colour is associated with a different cluster. C. Distance matrix representing the squared MMD between all lines from the inactivation screen, computed on a 10D learned latent space for a 2-second time window. D. 2D representation of the geometric relationships between lines, obtained using supervised UMAP, encoded by the distance matrix. The bar plot associated with each cluster represents the average variation of behaviour during the 2-second window in the six actions behaviour dictionary. The thickness of the lines linking the cluster is associated with the coupling between the clusters. E. The z-score average between data and generated sequences, as well as the overall lines, is divided by the standard deviation of these z-score distributions. We display only the 30 highest values. F. The 17 sequences of nodes with the highest frequency of occurrence for each of the eight clusters.

We apply the suffix tree o cluster different lines by utilizing internal nodes shared between several lines. Here, the advantage of using a suffix tree is the ability to compare sequences of different lengths while retaining the order of behaviors. In particular, we compare each node of the suffix tree, examining node overlaps across multiple lines in the suffix tree. To define a metric, we embed the behavioral sequences of each genetic line using the Vector Space Document (VSD) model. In this case, a document corresponds to a genetic line. We map the nodes from the common suffix tree to an *M* -dimensional space in the VSD model.

In the VSD model, each genetic line, *𝓁*, is considered as a vector in an M-dimensional term space. We construct these vectors by the term frequency-inverse document frequency (tf-idf) weighting scheme proposed in [62], [63]. It measures the relevance of a word (here, a node) according to its frequency in each document (line) and in a document collection (cluster of lines). The vector corresponding to a line *l* is given by

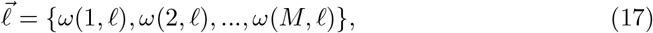

where *ω*(*i, 𝓁*) is the weight applied to each node *i* in document *𝓁*, defined by the tf-idf,

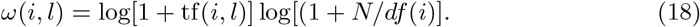

Here *N* is the number of lines, tf(*i, 𝓁*) is the frequency of the *ith* node in the line *𝓁*, and *df* (*i*) is the number of lines containing the *ith* node. The frequency is given by 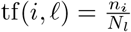, where *n*_*i*_ is the number of larvae that pass through this node *i* and *N*_*𝓁*_ is the number of total larvae in the line *𝓁*. We obtain a vector for each line, with a weight term for each node. We calculate a square distance matrix from these vectors containing the pairwise distances between the vectors of the genetic lines. We measure the distance between two vectors 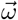 and 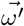 as the cosine similarity since it’s more robust to the variability in the number of larvae per line compared to the Euclidean distance,

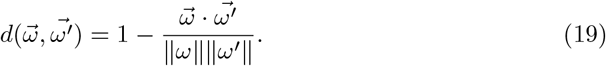

We apply hierarchical clustering to the distance matrix to group the genetic lines according to their behaviour sequences (Fig. 6B).

## Results

We applied our methods to generate behavioural phenotypic descriptions that are crucial for understanding both the global scale (across the entire screen) and the local scale (at the level of individual genotypes). We constructed a distance matrix encompassing all lines by analyzing the distribution of genetic lines within the latent space. To achieve a geometric perspective on the relationships between various genotypes at a large scale, we calculated the pairwise Maximum Mean Discrepancy (MMD) distance matrix. Our approach involved a two-step process: calculating the MMD distance matrix for all lines and then embedding the genotypes into a high-dimensional geometric space through multi-dimensional scaling. This space, another latent space, represents the probabilistic reactions to stimuli at the genetic line level. Using hierarchical clustering with Ward’s linkage method, we visualized this latent space (referenced as Fig. 6D), identifying five contiguous regions. For a conventional representation of the primary behaviour statistics, we calculated and compared the average behaviour histograms for lines within each region against the overall average behaviour histogram derived from this geometric framework. Additionally, we explored the interrelations among these regions using supervised Uniform Manifold Approximation and Projection (UMAP) for dimensionality reduction. This process resulted in each subregion being represented as a separate connected component in a 2D space. We also illustrated the total connectivity between each region, quantified by the sum of edge weights in the graph created through the UMAP algorithm, shown in dashed grey lines with widths proportional to the logarithm of the connectivity.

The scope of the screen, combined with the variety of genetic lines and behaviours, facilitates the categorization of larval dynamics into clusters and sequences of behaviours. By employing hierarchical clustering on the representation vectors of nodes within the suffix tree — which captures both the frequency of a sequence’s occurrence and the number of lines displaying it — we can depict the principal families of larval behaviours in response to this sensory stimulation paradigm (Fig. 6F). This approach reveals the larvae’s reactions to airflow natural stimuli, notably bending movements. A distinction emerges between response families characterized by the hunch (head retraction) and repetitive transitions between back movements and bends and those characterized by rapid escape involving running phases, stopping phases, and then swiftly resuming running and bending cycles. The latter represents the baseline behaviour of larvae placed on a 2D agar plate without a specific task, likely engaging in foraging behaviour.

Globally, the generative model allows for identifying behavioural sequences that are most likely unable to be described by a time-inhomogeneous Markov model (see Fig. 6E). These sequences, where back and hunch are often represented, are associated with the larvae avoidance manoeuvres. Numerous questions remain regarding how behavioural sequences are encoded and their neural implementation. Similarly, encoding the duration of these behaviour motifs needs to be investigated as some lines, for example, will exhibit 1 or 2 repetitions of the (back, bend) motifs while in others, for example .

Our principal finding comprises a catalogue of genetic lines exhibiting subtle behavioural modifications, as identified through statistical testing within the behavioural latent space and via the Bayesian generative model for action sequences. The genetic lines pinpointed by our methodologies are detailed in the Supplementary Information, with specific examples illustrated in Figure 7. We present the reference line alongside two instances of lines identified through the Maximum Mean Discrepancy (MMD) method and two others recognized by the generative Bayesian model. Both techniques have considerably broadened the spectrum of lines of interest by their capacity to pinpoint behavioural evolution that is not overtly manifested by significant changes in individual actions, either through their emergence or absence, as noted in [34]. Accordingly, each method uncovers two distinct sets of characteristics. Intricate patterns distinguish the lines newly identified by the MMD method (Fig. 7D) in the witness function landscape, indicating alterations across multiple behavioural domains. Not all these modifications align with the behaviours defined by the discrete action dictionary.

**Fig 7.**
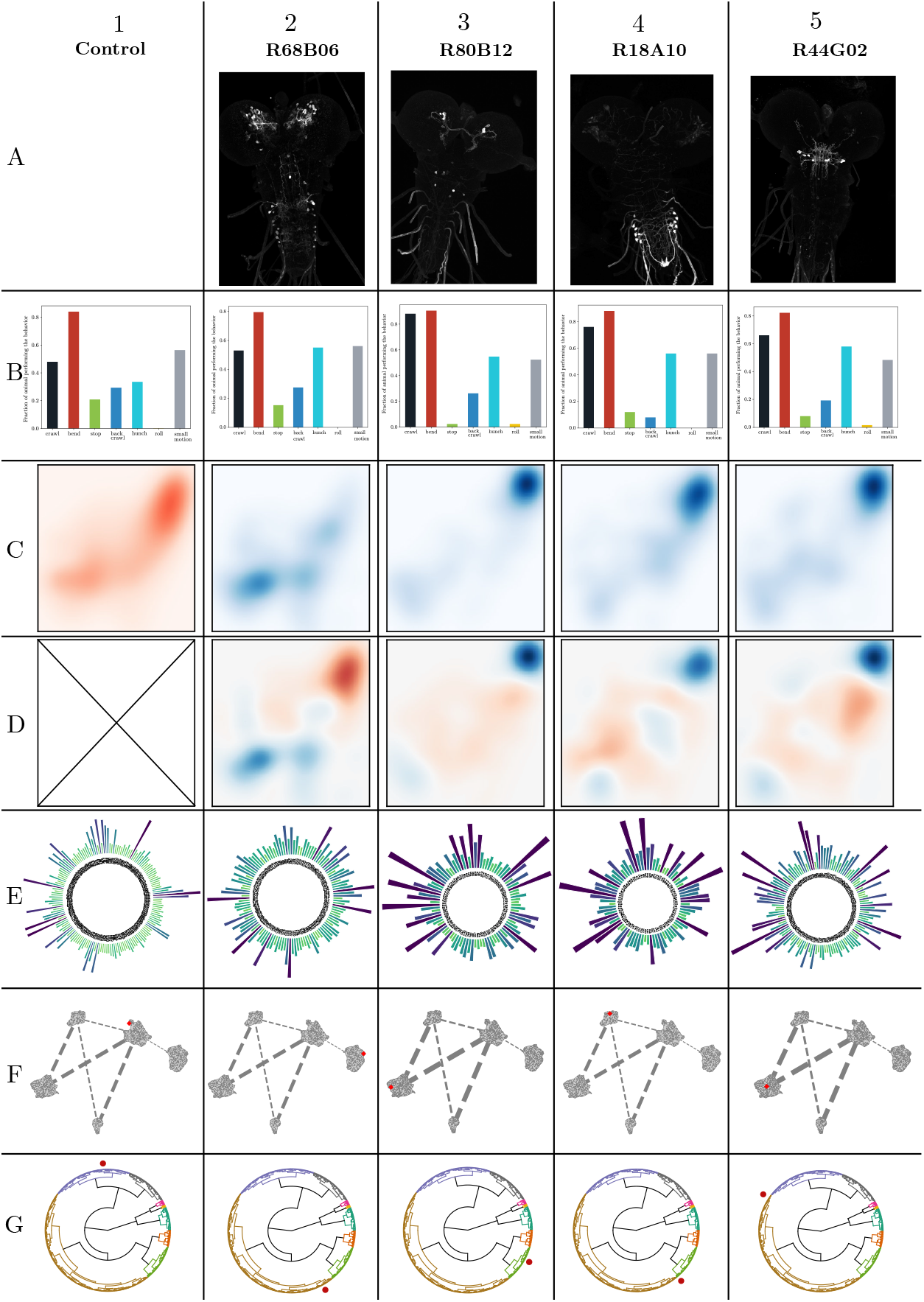
Samples of genetic lines of interest, ”Hits”, with their characterisation. These lines lead to subtle modifications of behaviour and were not detected by previous approaches. We present four new hits: two hits associated with complex alterations of the learned latent space and two lines associated with strong sequence deviations from the generative model and the reference. The columns correspond to 1. control line *attP* 2, 2. *R*68*B*06, 3. *R*57*F* 07, 4. *R*18*A*10, 5. *R*38*H*09. Row A: Light microscopy images of larval brains expressing the selected GAL4 line. Note that there is no picture for *attP* 2 as it is the reference and thus labels no neurons. Row B: Fraction of behaviour during 2 seconds after the stimuli projected onto the six action dictionary. Row C: Latent normalized distribution of behaviour of the lines, during 2 seconds after the stimuli, with in red the distribution of the reference and in blue the distribution of the hits line. Row D: Witness function between latent distributions highlighting the main sources of behavioural differences between the lines. Note the complex patterns in the latent space, showing these hits don’t stem from simple variations in one action. Row E: z-score of sequences of three actions between generative and experiment sequences. Row F: Position of the reference and hits lines in the 2D representation of the geometric relationships between lines encoded by the distance matrix (shown in Fig 6D,E). Row G: Position of the reference and hit lines in the hierarchical clustering tree (shown here in circular form).

The new lines identified by the generative probabilistic model are characterized by longer-term effects on action sequences, as illustrated in Figure 7E (these lines are listed in Supplementary Table 5). Our findings reveal that these lines display variations in the global proportions of specific sequences of three actions despite having average probabilities of individual actions comparable to the reference line. For sequences beyond three actions, the statistical significance of the findings could not be guaranteed across all lines. The newly detected lines were discovered across a broad range of the screen in clusters defined by either the suffix tree representation or the MMD-based distance matrix, as shown in Figure 7F–G.

Our methods successfully identified nearly all the hits previously reported by Masson et al. (2020) [34] as strong hits (Table 4). However, some hits (listed in Supplementary Table 6) are no longer classified as such according to the more stringent criteria of our two new approaches. There are several factors contributing to their reclassification. In many instances, transitioning to a definition of behaviours within a continuous latent space—and away from the discrete categorization of behaviour—eliminates strict boundaries, leading to a loss of significance under the current methodology. It is important to note that different conceptualizations of behaviour may yield varied criteria for significance. An increase in the sample size of larvae from these lines will be crucial in determining whether they still qualify as hits under these revised definitions.

In this study, we extended the behavioural paradigms to include a subset of lines (referenced in Table 7) to examine their behavioural responses to varying levels of air puff intensity. As previously reported, larvae exhibit different behaviours in response to lower stimulus intensities, such as fewer hunches, bends, and backups, and an increase in stops and crawls [34] (see Fig. 2). Figure 8 presents two lines that demonstrate distinct phenotypic variations in their modulation of behavioural responses to different air puff intensities. The first line, R68B05 (shown in column 2 of Fig. 8), shows a phenotypic difference between the intensities—displaying more pronounced hunching in response to high intensity and less at low intensity compared to the reference. This results in a greater disparity in hunch response probabilities between low and high stimuli compared to the control, suggesting that neurons in this line may play a role in modulating control and maintaining a stable range of behavioural responses regardless of stimulus intensity. The second line, R20F11 (illustrated in Fig. 8, column 2), exhibits a consistent phenotype across both intensity levels, indicating an absence of behavioural modulation based on stimulus intensity (Fig. 8A). Neurons in these lines might thus be implicated in encoding stimulus intensity and/or regulating the behavioural response in a stimulus-intensity-dependent manner. Our new methods further supports this phenotypic distinction; the witness function uncovers a significant difference in response to high versus low intensity for one protocol compared to the reference, whereas the other displays minimal variation. We can subsequently locate the positions of these two protocols within the latent space and the suffix tree of action sequences å.

**Fig 8.**
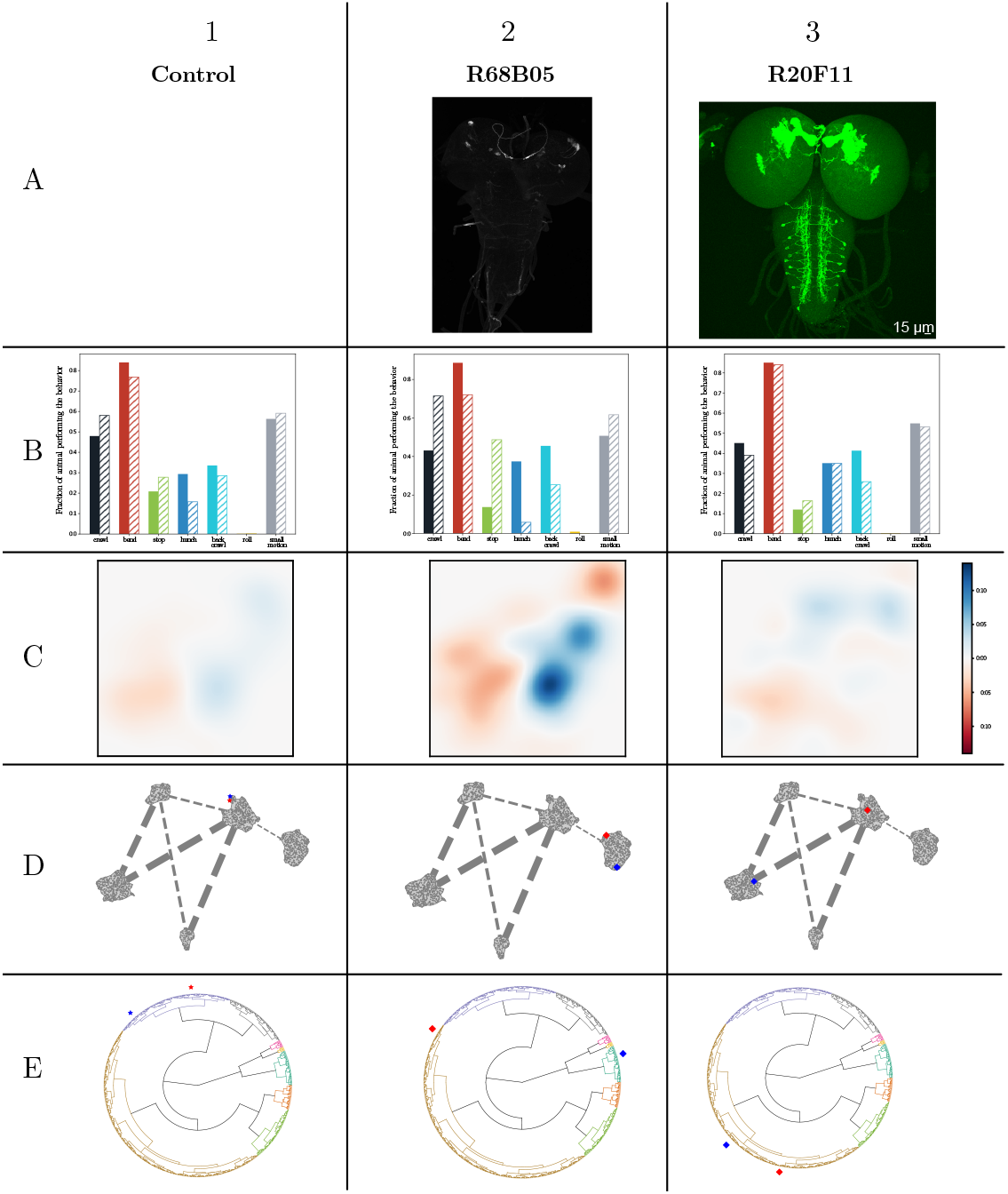
Two genetic lines of interest, each subjected to two different stimulus intensities: high intensity as previously illustrated, and low intensity, involving a less powerful air puff. We provide characterizations of each line and protocol. The columns correspond to (1) the control line, (2) *R*68*B*05 and (3) *R*20*F* 11. Row A displays light microscopy images of larval brains expressing the selected GAL4 lines. In Row B, the fraction of behaviour during the 2 seconds following the stimuli is projected onto the six-action dictionary, with high intensity in plain colour and low intensity in dashed lines. Row C, the witness function between latent distributions, highlights the main sources of behavioural differences between the two protocols for the control and the two lines. Row D, the position of high intensity in red and low intensity in blue in the 2D representation of the geometric relationships between lines, encoded by the distance matrix (as shown in Fig. 6B,C). In Row E, the position of high intensity in red and low intensity in blue is displayed in the hierarchical clustering tree (presented here in circular form)

## Discussion

The swift progress in large-scale behavioural studies, complemented by neural manipulations and recordings, paves the way for establishing causal connections between behaviour and its neural underpinnings. Although various statistical methods can identify immediate and pronounced deviations, detecting subtler variations remains challenging. These minor deviations may stem from nuanced behavioural changes that are difficult to detect or from modulations happening across challenging-to-capture timescales.

Our ability to detect such nuanced modulations arises from our decision to simplify behavioural quantification into two distinct measurements. In the first approach, we linked behavioural features with the mechanical consistency of larval movements. We then reduce the dynamics of population action over a set time scale, representing it as a distribution within a learned latent space. Thanks to the low dimensionality of our latent space, we employed robust kernel-based statistical tests. As a result, our detection of subtle behavioural responses outperformed those based on dictionary-based projections. It allowed the extension of behaviour detection at the frontier of ambiguous actions to the expert eye. Hence, the complex patterns exhibited by the witness function (examples shown in Fig. 7C) for these lines of interest often include the boundaries between previously described discrete behaviours (see Fig. 3).

We developed a streamlined self-supervised method to encode actions in a continuous latent space, employing a compact neural network. This approach is adaptable to various architectures, enabling the creation of a meaningful, continuous latent space. It can also be integrated with different interpretations of what constitutes a behaviour or an action. In that sense, approaches looking for underlying behavioural structures in the spatiotemporal dynamics of postural movement data, [21, 24, 26], in the latent structure of animal motion prediction [23], or in continuous latent spaces compressing raw video of behaviour [27] could be directly patched into our procedure. In this context, the primary limitation arises from statistical testing. As the dimensionality of the latent space increases, so does the risk of anomalies [64–68], a phenomenon known as the curse of dimensionality.

The approach could be extended to larger timescales without retraining but simply by encoding more significant epochs into multiple points, then defining a distribution on the latent space and comparing conditions using MMD-based statistical testing. However, since the ordering of these points would not be represented, long-time window encoding will lose resolution in the temporal sequences of action and, thus, part of the relevant information. Although Maximum Mean Discrepancy (MMD) exhibits some resilience against the escalation in data dimensions [57, 69], the finite number of genetic lines and corresponding larvae implies that a lower-dimensional latent space would enhance the statistical significance of the analysis.

The neural circuitry underlying individual actions and sequences of actions remains partially understood and modelled. There are ongoing debates [70–77] regarding the mechanisms of action initiation, temporal stability, and the transition into new actions. These discussions revolve around whether these processes are localized in specialized centres with centralized competition or are distributed throughout the nervous system. Similarly, there is active debate about the control mechanisms governing the sequence of actions, with models such as chains of disinhibitory loops [10], parallel queuing [78], ramp-to-threshold [79, 80], and synaptic chains [81, 82] under consideration. Our second approach focuses on uncovering complex correlations within the structure of action sequences at the population level. Although behaviour alone may not conclusively pinpoint the neural mechanisms responsible for generating sequences, complexities observable at the population scale—like non-Markovian characteristics or high-order correlations—may offer clues about the neurons that orchestrate these dynamics in sequence generation.

In our second approach, we utilize the structured framework of a tractable probabilistic generative model to explore complexities in action sequences. This model is a foundation for contrasting a group of larvae against a corresponding constrained reference model, eliminating the need for an external reference line for comparison. Our method is adept at identifying complex temporal variations in sequences at the population level, thanks to analysing higher-order correlations within these sequences and comparing them against the constraints of the generative model. In this setup, the future state of an individual is determined solely by its current state, which is in line with a Markov model. The model also incorporates variability through a potentially time-varying effective action rate. This approach has enabled the identification of genetic lines where the time-evolving probabilities of actions align with experimental data, albeit with limitations. For instance, it does not account for the frequencies of sequences of three actions, among others. Thus, these identified ”hit” lines reveal limitations in capturing certain aspects of action sequence generation when simplifying the dynamics to an inhomogeneous Markovian Poisson model.

The intricacies of behaviour and its connection to neural computations, whether in specialized circuits or distributed across the nervous system, cannot be fully understood through a single method or confined to a particular time scale. However, we can identify meaningful behavioural characteristics by integrating multiple methods that utilize both local and global data at the level of individual animals and across populations and by spanning various time scales. These characteristics can then be compiled into behavioural identity documents for individual neurons and neuronal clusters (refer to Fig. 7). By merging these detailed profiles with connectome data [4] and neural recordings, we can hasten the discovery of circuits responsible for decision-making and the subtle nuances in their output.

## Supporting information

### Latent space

#### New behaviours

We tested the capacity of the latent space to represent new actions beyond the six classical ones [10, 34]. We show two examples of possible new action categories in Figure 9, namely C-shape (in A) and head-tail (in B). In the former, the larva takes the shape of a C with variable time spent in that state. It is observed, for example, prior to rolling. The larva exhibits hunch-like motion in the latter with rapid head and tail retraction. It is observed, for example, following air flow puffs.

**Fig 9.**
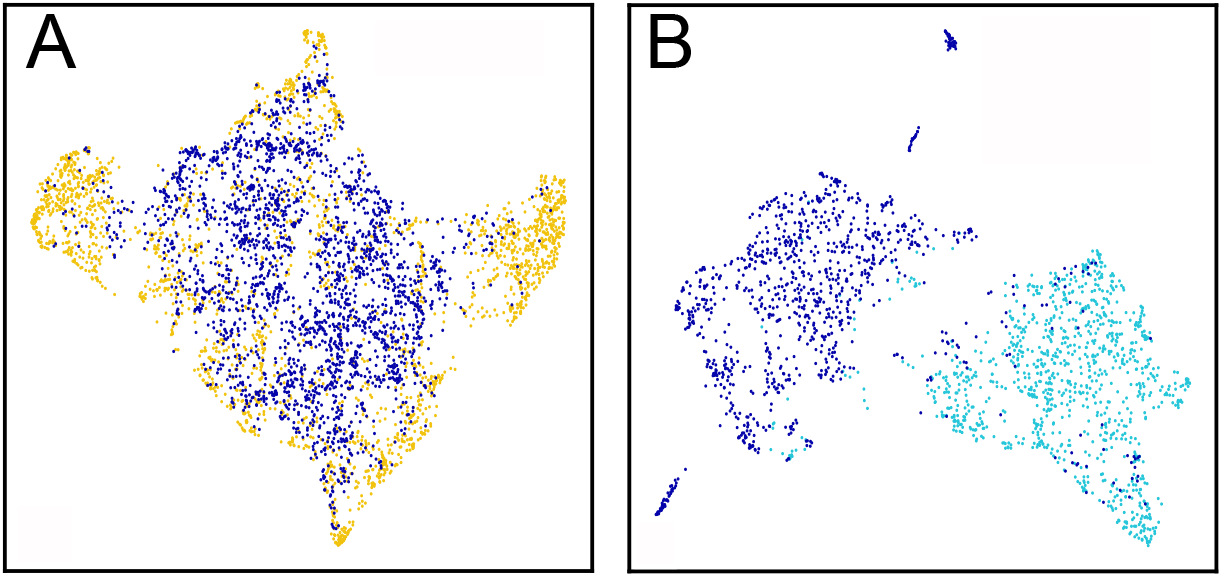
Latent representations of C-shape (A)and Head-tail (B). (A) C-shapes: deep blue, rolls: yellow. (B) Head-tails: deep blue, backs: light blue.

#### Exploring the Latent Space

Beyond the usual dictionary of larval actions and behaviour, the latent representation can be explored without labels.

#### Clustering the latent space

While clustering is not necessary for our analysis approaches, discrete behaviour description can be instrumental in describing larva dynamics.

We used the persistence-based clustering algorithm ToMATo [83]. The algorithm requires an estimate of the density at the data points and the pairwise distances matrix between them to perform the clustering. The clustering joins a mode-seeking phase based on a graph-based hill-climbing scheme and a topological persistence merging phase in the density map. Clustering was performed on the combined training and validation dataset.

In ToMATo [83], the number of clusters is controlled by a merging threshold, the minimum prominence a local peak must attain to be considered significant. A common practice to define the number of clusters is to use the gap statistic [84]. Instead of setting the number of clusters, we designed a graphical interface to examine the hierarchy dynamically. Interestingly, as in [3], one of the clusters identified through this procedure captures an anomaly in the larva tracking where the head and the tail are suddenly swapped.

#### Interface to navigate the latent space

We developed a software tool which allows for interaction and visualization of the cluster hierarchy, a visualisation of the 2D projection of the latent space, and generation of video data representing the samples in each cluster (see Fig. 10).

**Fig 10.**
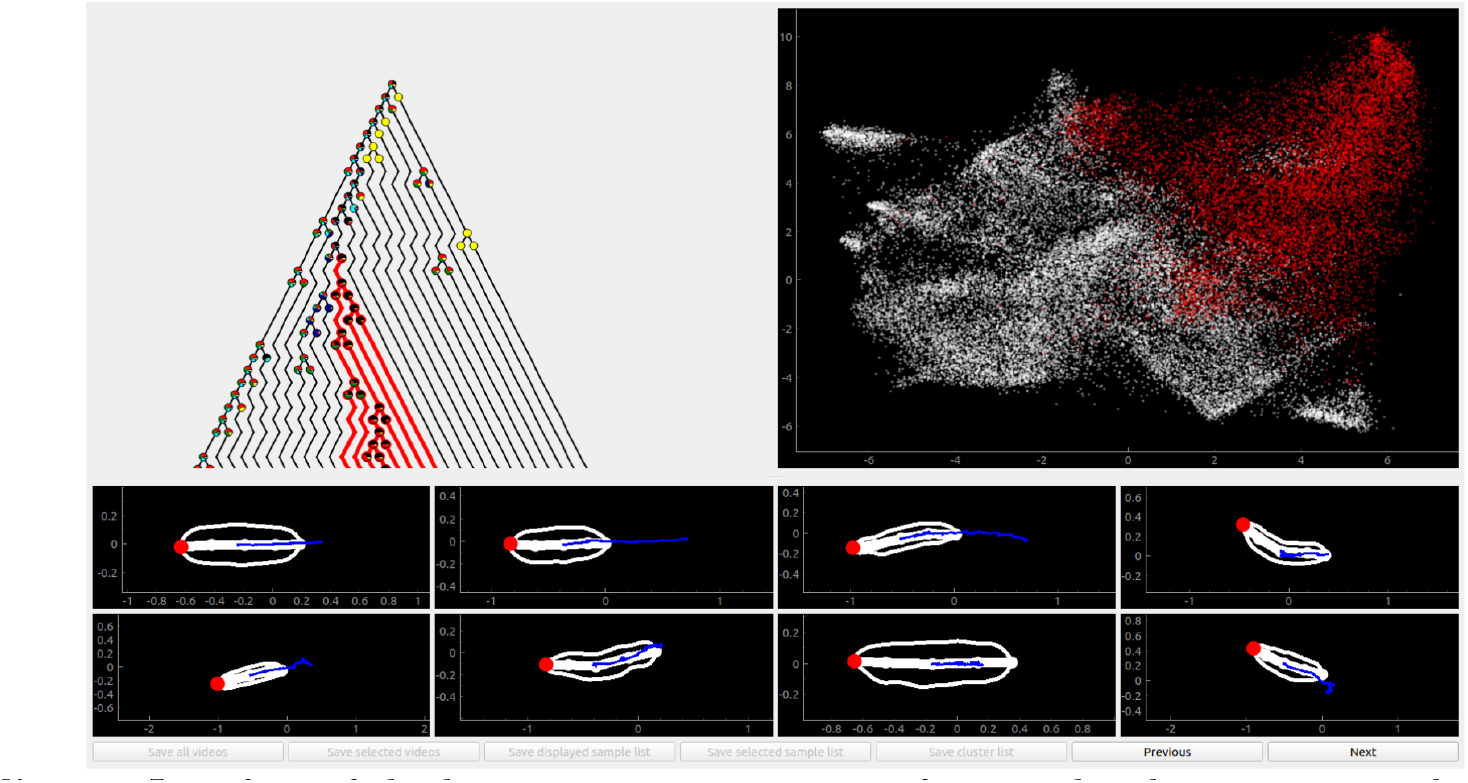
Interface of the latent space navigation software: the clustering tree, the portion of the latent space selected, and examples of larva dynamical actions belonging to this cluster. Supplementary Videos 2. provides a video showing interaction with the software.

By interacting with the tree on the top left, we can choose a particular cluster to visualize. The cluster node and its children are then highlighted in the tree, while the corresponding data points are highlighted in the 2D projection on the top right. Samples from the cluster are displayed at the bottom of the interface. Various display settings can be used, such as the display of the outlines or the midlines of the recorded larva contours. The larva’s head is highlighted in red, while the trace of its midpoint is plotted in blue. Finally, the depth of the cluster tree can be varied.

To declutter the tree view, the user can interactively fold a cluster, hiding all of its children from view, if they consider the distinctions between the different children clusters irrelevant. Note that these merges need not be consistent with the merging criterion of ToMATo, leaving the researcher with all the freedom to merge clusters, with the limitation that cluster merges must respect the tree structure.

### Screen scale cluster definition

The screen can be used to define relevant cluster numbers. After computing the complete cluster hierarchy and associating each genotype to all clusters, we can prune the hierarchy to ensure that all clusters have at least one genotype belonging to them that is different from other clusters.

### Generative model

To generate the numerical sequences of actions, we rewrite the likelihood shown in Eq. 15 as

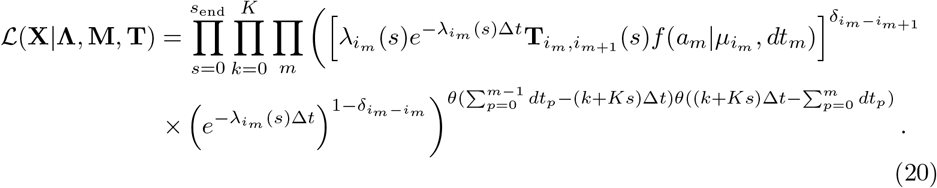

In this form,

#### Algorithm 1

MCMC to generate behavioural sequences

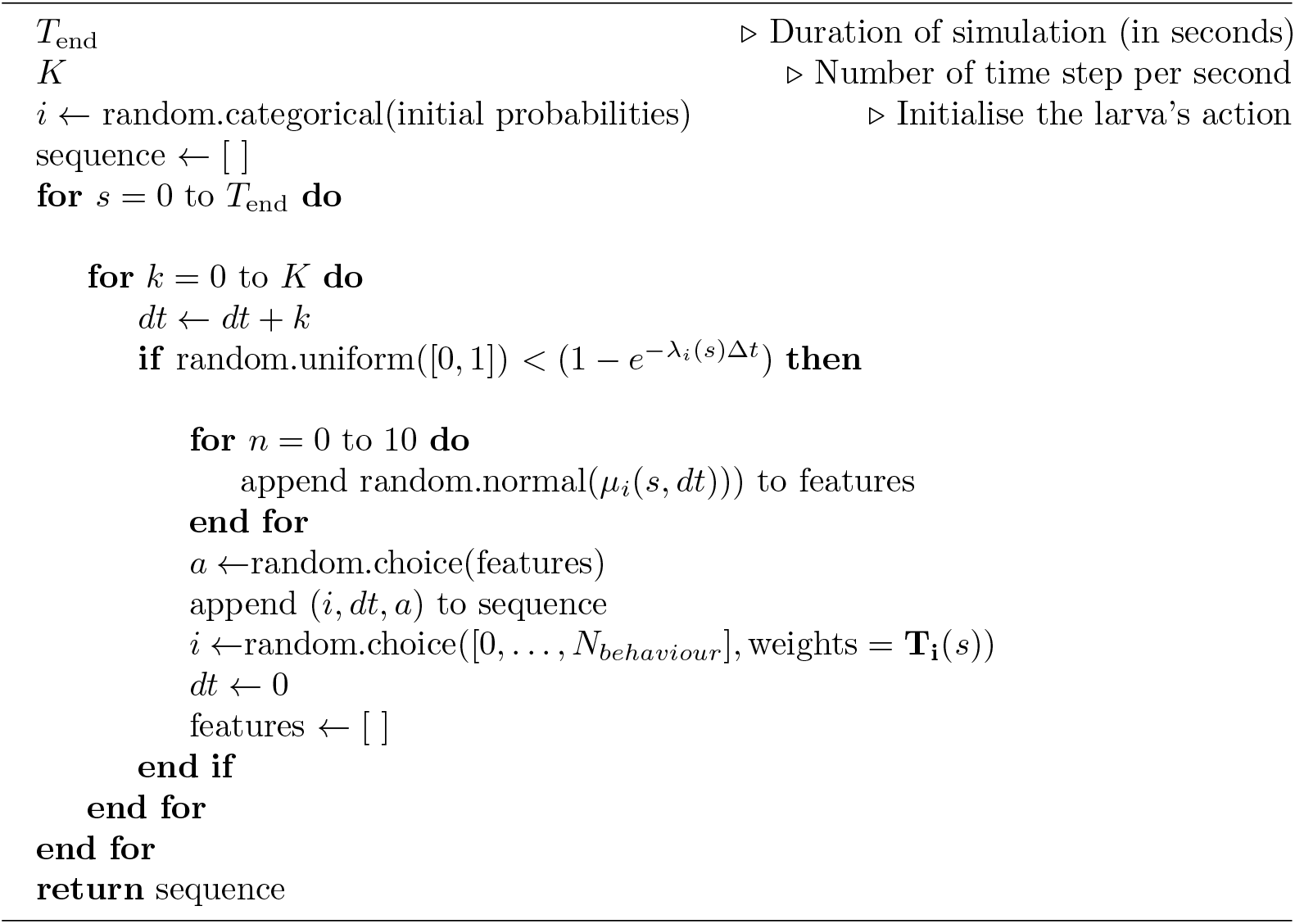

#### Likelihood ratio test

We employ a likelihood ratio test that compares the lines to the reference line to detect behavioural modifications of genetic lines. The test statistic is given by

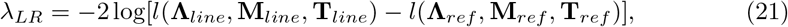

where *l* is the logarithm of the maximized likelihood function *L*. This test is made possible as the generative model provides a tractable likelihood.

## Tables

- Table 1. List of lines studied.
- Table 2. A table presenting the estimated square of the latent space Maximum Mean Discrepancy (MMD) with the corresponding bootstraped p-values for all lines, complemented by a calculation of the distance between generative and experimental sequences.
- Table 3.1 and Table 3.2. Z scores comparing action sequences from the generative model to experimental recordings obtained respectively during the stimulus and at all times.
- Table 4. A table presenting the estimated square of the latent space Maximum Mean Discrepancy (MMD) and the distance between generative and experimental sequences for lines previously identified as hits, as cited in [34], and confirmed as such with this analysis.
- Table 5. A table providing the list of new hits detected by the two new approaches.
- Table 6. A table providing the few genetic lines that were detected as hits in [34] and that no longer are hits with this analysis.
- Table 7. A table detailing the estimated square of the latent space Maximum Mean Discrepancy (MMD) with the corresponding bootstraped p-values, complemented by a calculation of the distance between generative and experimental sequences for lines with low and high intensity of the stimulus.

## Acknowledgements

This study was funded by the INCEPTION project (PIA/ANR-16-CONV-0005), the *“Investissements d’avenir”* program managed by *Agence Nationale de la Recherche*, reference ANR-19-P3IA-0001 (PRAIRIE 3IA Institute) to J.B.M, C.L.V, C.B & A.B, and *Agence Nationale de la Recherche* ANR-20-CE45-0021 to C.L.V. and A.B. This work was supported by ANR PIA funding: ANR-20-IDEES-0002 (T.J), Agence Nationale de la Recherche (ANR-17-CE37-0019-01) (T.J), ANR-NEUROMOD (ANR-22-CE37-0027) (T.J). This project has also received funding from the European Union’s Horizon 2020 research and innovation program under the Marie Sklodowska-Curie grant agreement No 798050 (T.J & J.B.M). L.M was supported by the Amgen Scholars program.

