## Supplementary figures and images for "Statistical signature of subtle behavioural changes in large-scale behavioural assays"

### Supplemental Video 1

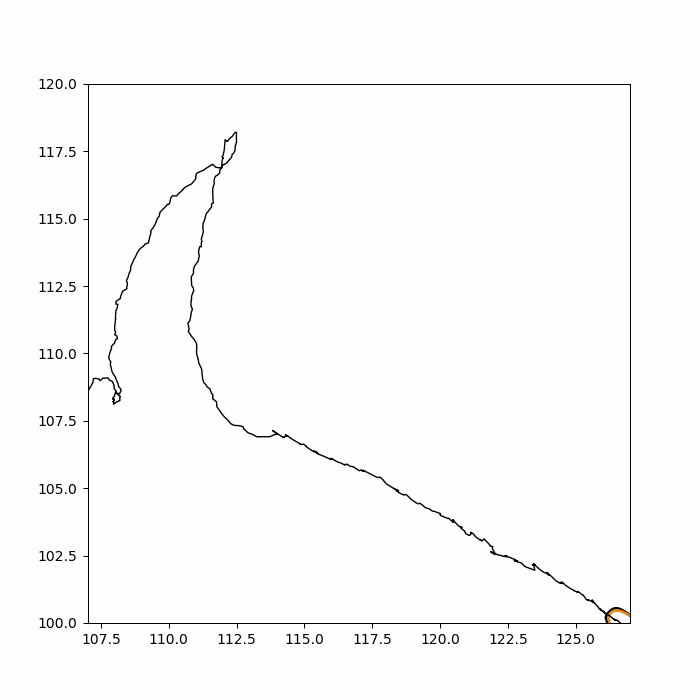
